# When fibers crowd: Mesh burden and myofibroblast phenotypes in women with vaginal mesh exposure

**DOI:** 10.64898/2026.11.13.766516

**Authors:** Malini Harinath, Marrisa A. Therriault, Jordan Head, Misha Patel, Katrina Knight, Amanda Artsen, Pamela A. Moalli

## Abstract

Polypropylene mesh improves the durability of pelvic organ prolapse (POP) repair but can cause complications, most commonly vaginal mesh exposure. Preclinical animal studies demonstrate that mesh deformation increases local mesh burden and is associated with maladaptive remodeling and macrophage-to-myofibroblast transition (MMT). Whether a similar cellular response occurs in women remains unknown. We examined mesh–vagina tissue complexes excised from N=17 women with vaginal mesh exposure following sacrocolpopexy. Specimens were stratified into quartiles of mesh burden and evaluated for macrophage density, fibroblast-to-myofibroblast transition (FMT)- and MMT-associated phenotypes, cellular localization relative to mesh fibers, and collagen content and organization. We also interrogated our previously published proteomic atlas, generated from the same institutional mesh excision biorepository, to compare molecular signatures in vaginal tissue from women with mesh exposure, mesh-associated pain, or well-integrated mesh and controls without mesh. Macrophage density was greater in the highest mesh burden quartile than in the two lowest quartiles. Across all quartiles, approximately 96–99% of myofibroblasts co-expressed macrophage markers, consistent with an MMT-associated phenotype, and localized near the mesh– tissue interface. MMT-associated cell density increased numerically with mesh burden, although differences were not statistically significant. Collagen content relative to mesh area was lower in the higher burden quartiles, whereas collagen as a percentage of tissue area did not differ. Proteomic analysis identified alterations in macrophage-associated and matrix-remodeling proteins, with prominent changes in exposure specimens. These findings identify a predominant MMT-associated myofibroblast phenotype in women with vaginal mesh exposure and link greater mesh burden to heightened macrophage accumulation and reduced collagen relative to implanted material. MMT-associated pathways warrant further investigation as potential therapeutic targets for maladaptive mesh remodeling.

## 1.0 Introduction

Pelvic organ prolapse (POP) is a common disease in aging women in which the pelvic organs descend into the vagina because of weakening or injury to the supportive pelvic connective tissues and muscles. Nearly half of women over age 50 experience symptoms of POP with the risk of surgery estimated at 13% of all US women by age 80 [1, 2]. Polypropylene mesh (PPM) provides durable mechanical support for prolapse repair, particularly in sacrocolpopexy, but its benefits are tempered by mesh-related complications, including pain associated with fibrosis and mesh exposure through the vaginal epithelium [3, 4]. Understanding the mechanisms that distinguish successful mesh integration from maladaptive remodeling is essential to improving the safety and durability of these repairs [5, 6].

Macrophages are central regulators of the foreign body response, coordinating inflammatory signaling, fibroblast activation, and extracellular matrix turnover at the mesh–tissue interface [7–10]. Although a persistent foreign body response is expected around a permanent implant, excessive or dysregulated inflammation can compromise tissue integration [11–14]. Myofibroblasts also contribute to pathologic remodeling through their high matrix production and contractile activity resulting in fibrosis and pain [15]. These cells are conventionally considered to arise from activated fibroblasts in the fibroblast to myofibroblast transformation (FMT); however, recent evidence suggests that macrophages can also acquire myofibroblast features through macrophage-to-myofibroblast transition (MMT) [16–18]. In our non-human primate model of sacrocolpopexy, mesh was implanted in configurations that progressed from a low (optimal for mesh integration) to high (reproduces complications) mesh burden. Interestingly, mesh burden scaled with the proportion of myofibroblasts co-expressing macrophage markers, or MMTs [19, 20]. These cells localized near mesh fibers and when abundant were associated with a less mature matrix, decreased collagen density, and increased TGF-β1 [19]. Together, these findings suggest that MMT may contribute to maladaptive remodeling in regions of high mesh burden.

Whether a similar response occurs in women remains unknown. Human mesh explant studies have demonstrated persistent macrophage-dominated inflammation, including pro-inflammatory macrophage phenotypes, with more limited changes in T-cell recruitment in women with complications [13, 21]. However, the relationship between local mesh burden, myofibroblast phenotype, and matrix remodeling has not been established in human vagina. Moreover, although our primate model permits controlled investigation of mesh geometry and loading, human specimens capture the heterogeneity of patients, implantation histories, and long-term tissue responses encountered clinically. A recently published proteomic atlas of mesh–vagina specimens from women with pain and/or exposure and women without mesh complications provides an additional opportunity to investigate molecular pathways associated with adverse remodeling [23].

In this study, we investigated whether increasing local mesh burden is associated with an MMT-associated cellular phenotype and altered collagen content in women with complications, primarily exposure, following mesh-augmented pelvic organ prolapse repair surgery. We hypothesized that greater mesh burden would be associated with an increased proportion of myofibroblasts co-expressing macrophage markers, consistent with the response observed in our primate model. To test this hypothesis, we stratified mesh–vagina explants by quartiles of mesh burden (fiber per area) and quantified collagen content and myofibroblast populations according to macrophage marker expression. We then interrogated the previously published human proteomic dataset to determine whether specimens from women with complications showed enrichment of proteins and pathways implicated in MMT compared with specimens from women without complications. Together, these complementary analyses examine the clinical relevance of our preclinical findings and the potential contribution of MMT to adverse remodeling following mesh-augmented prolapse repair.

## 2.0 Methods

### 2.1 Sample selection and inclusion criteria

Samples via the Magee Women’s Mesh Biorepository, a large, IRB-approved human urogynecologic mesh biorepository at the Magee Womens Research Institute (IRB = CR19080175-010). All patients undergoing excision of vaginal mesh placed at the index surgery for prolapse or urinary incontinence are offered enrollment. Biologic samples, including excised vaginal-mesh complexes, are stored in an ultra low-temperature freezer until use. Demographic and medical data are prospectively abstracted and Pelvic Floor Distress Inventory-20 and visual analog scales for pelvic pain are collected. For this study, samples were included if they were a sacrocolpopexy or vaginal mesh with a complication (exposure, pain, mesh buckle) and sections included both mesh fibers and adequate tissue for analysis. Out of 38 participants that met the criteria, 26 had mesh-tissue complexes available for histological analysis.

### 2.2 Cryosectioning and slide preparation

All mesh-tissue complexes were preserved in OCT and frozen in an ultra low-temperature freezer immediately following mesh excision and sample procurement. Sample cross-sections were cut into 7 µm thick sections using a Leica Cryostat (Nussloch, Germany) and placed on negatively charged slides.

### 2.3 Hematoxylin and Eosin

Frozen sections were fixed using 4% paraformaldehyde at 4C for 1 hour. Slides were then washed with 1x PBS and placed in Harris hematoxylin (Fischer Scientific, Waltham, MA, USA) at room temperature for 8 minutes. Slides were then rinsed with tap water followed by Scotts tap water before being placed in Eosin Y (Fisher Scientific, Waltham, MA, USA) at room temperature for 1 minute. Slides were then dehydrated using an ethanol gradient (95-100%) and xylenes before being mounted with Clear Vue mount (Fischer Scientific, Waltham, MA, USA). The slides were imaged using the Motic Easy Scan Pro Digital slide scanner at a 10x magnification to ensure adequate cell and tissue presence for histologic analysis.

### 2.4 Cell and area quantification

Total tissue area, total mesh area, and total area of tissue without mesh fibers were outlined and quantified. Mesh burden was quantified by calculating the percentage of tissue area occupied by mesh [20] and separated into four quartiles resulting in N=4-5 per group for further analysis through the rest of the outcomes.

### 2.5 Trichrome staining for total collagen content

OCT preserved, cryosectioned samples were fixed using acetone at -20C for 20 minutes and washed using 1x PBS and 95% ethanol. The tissue sections were placed in heated Bouins’ solution followed by Weigert’s iron hematoxylin solution for nuclear staining and Trichrome AB solution (Sigma Aldrich, Darmstadt, Germany). Decoloring was performed with 1% glacial acetic acid solution, and dehydrated using 95% ethanol, 100% ethanol, and xylenes prior to mounting using ClearVUE. The samples were imaged using the Motic Easy Scan Pro Digital slide scanner and the adventitial layer and mesh fibers were outlined. Collagen percentage was quantified using QuPath as the the proportion of blue pixels in each sample.

### 2.4 Immunofluorescence labeling

OCT mounted slides were fixed using 4% paraformaldehyde for one hour and then washed thrice using a 1x PBS. Sections were then covered with a 10 mM copper sulfate/50 mM ammonium acetate solution and incubated at 37C for 20 minutes. Following this, the samples were washed thrice using 1x PBS and incubated with 4% donkey serum blocking solution and 0.3% Triton-X in 1x tris buffered saline (TBS) for 30 minutes at room temperature. The sections were then incubated with primary antibodies to alpha smooth muscle actin (1:100, Abcam, Cambridge, UK, ab21027), vimentin (1:100, Novus Biologicals, Ontario, NB300-223SS), and CD68 (1:100, Abcam, Cambridge, UK, ab955) in 2% donkey serum overnight at 4C. The following day, the primary solution was washed off the samples using 1x TBS solution mixed with 0.1% Tween 20. A solution of secondary antibodies made in 4% donkey serum was applied at a 1:200 concentration, and the sections were incubated for two hours at room temperature. Following this incubation, sections were rinsed and washed with three 15-minute cycles of 1x TBS with 0.1% Tween 20 and then three 5-minute cycles of 1x PBS. Slides were mounted using Vectashield with DAPI (Vector Labs, Newark, CA). The samples were imaged at 10x magnification using a Nikon 90i microscope (Nikon USA, Melville, NY). The adventitial layer was outlined and analyzed using QuPath (QuPath 0.6.1). Mesh area was excluded from tissue area measurements to determine sample ROI. Only cells that were labelled with DAPI (nuclear) were considered in the analysis. Co-labelling of DAPI with vimentin, αSMA, and/or CD68 was used to classify the responding cells into the following cell populations: fibroblast (Vimentin+), fibroblast-to-myofibroblast transitioned cells (FMT, Vimentin+ αSMA+), macrophages (CD68+), and macrophage-to-myofibroblast transitioned cells (MMT, Vimentin+ αSMA+ CD68+).

### 2.6 Picrosirius red

OCT preserved, cryosectioned samples were fixed using acetone at -20C and washed using deionized water. The samples were then placed in a previously described picrosirius red solution [19] for an hour. Slides were decolorized using acetic acid water made using a 1:2 ratio of glacial acetic water to deionized water. The samples were placed in 95% ethanol, 100% ethanol, and xylenes prior to coverslips being applied using ClearVUE mounting media. The tissue sections were imaged on the Nikon ECLIPSE 9E microscope using polarized light at 10x magnification. Thick and thin collagen fibers in the vaginal tissue were distinguished and quantified using a previously published MATLAB code that measures amounts of red and green pixels [20].

### 2.8 Proteomic analysis

To identify the protein changes associated with mesh placement, analysis was performed on a previously published proteomics data set consisting of 40 patient samples. Mesh-exposure (n=10), mesh-pain (n=10), and well-integrated mesh groups (n=10) were individually compared against vaginal control biopsies (n=10) and expressed as log2fold change. Samples were lysed, digested, and normalized to 50µg total protein and nano-scale liquid chromatographic tandem mass spectrometry performed as previously described [5]. Forty ECM-related proteins were selected a priori for a hypothesis-driven differentially expressed protein analysis.

### 2.9 Statistical Testing

A sample size was calculated based on previously published data assessing the MMT response with increase mesh burden in a primate model of prolapse repair. Based on the observed difference in MMT cells out of responding fibroblasts in samples with high and low mesh burden in a study previously described [19], a sample size of 4 per group was sufficient to detect differences with a power of 0.9 and an alpha less than 0.05 using a two-tailed test for significance. All statistical analysis was conducted using GraphPad Prism 10 (GraphPad Software LLC, Boston, MA). Differences in demographic categorical variables between groups were assessed using Fishers exact test. Normality was tested using Shapiro-Wilk test. A one-way ANOVA was performed on normally distributed data and post-hoc comparisons were made using Tukey’s multiple comparisons with a single pooled variance if p<0.05. For non-parametric data or normally distributed with unequal variance between groups, Kruskal Wallis was used. A two-way ANOVA was run on the cell distances from nearest mesh fiber with Tukey’s multiple comparisons with a single pooled variance used to determine statistical significance between groups and distances.Significant differences were determined with an α < 0.05 and a 95% confidence interval.

## Results

### 3.1 Patient demographics

Twenty-six total patient mesh-vagina complexes from the biorepository were screened, and 9 samples were ruled out due to inadequate tissue present. Of the 26 total participants 26% underwent a transvaginal mesh repair and 74% underwent sacrocolpopexy. The primary indication of mesh removal was exposure (N=14), while N=1 indicated only pain, and N=2 had a palpable painful mesh buckle. Patients were divided into quartiles based on mesh burden. Within the quartiles, there was no significant difference in the mean age in years (p = 0.2) or BMI (p=0.3, range 23-28). All patients in the study were white, with no participants identifying as Hispanic/LatinX. There was no significant difference in smoking or diabetes status (p>0.05). There was no difference in the median gravidity or parity of the patients in each quartile (p>0.05). All patients in the study were postmenopausal. Seventy-one percent (n=12) of the patients in the study used some form of hormone replacement therapy. There was no significant difference in the mean duration of mesh implantation between quartiles (p>0.05, **Table 1**).

**Table 1.**
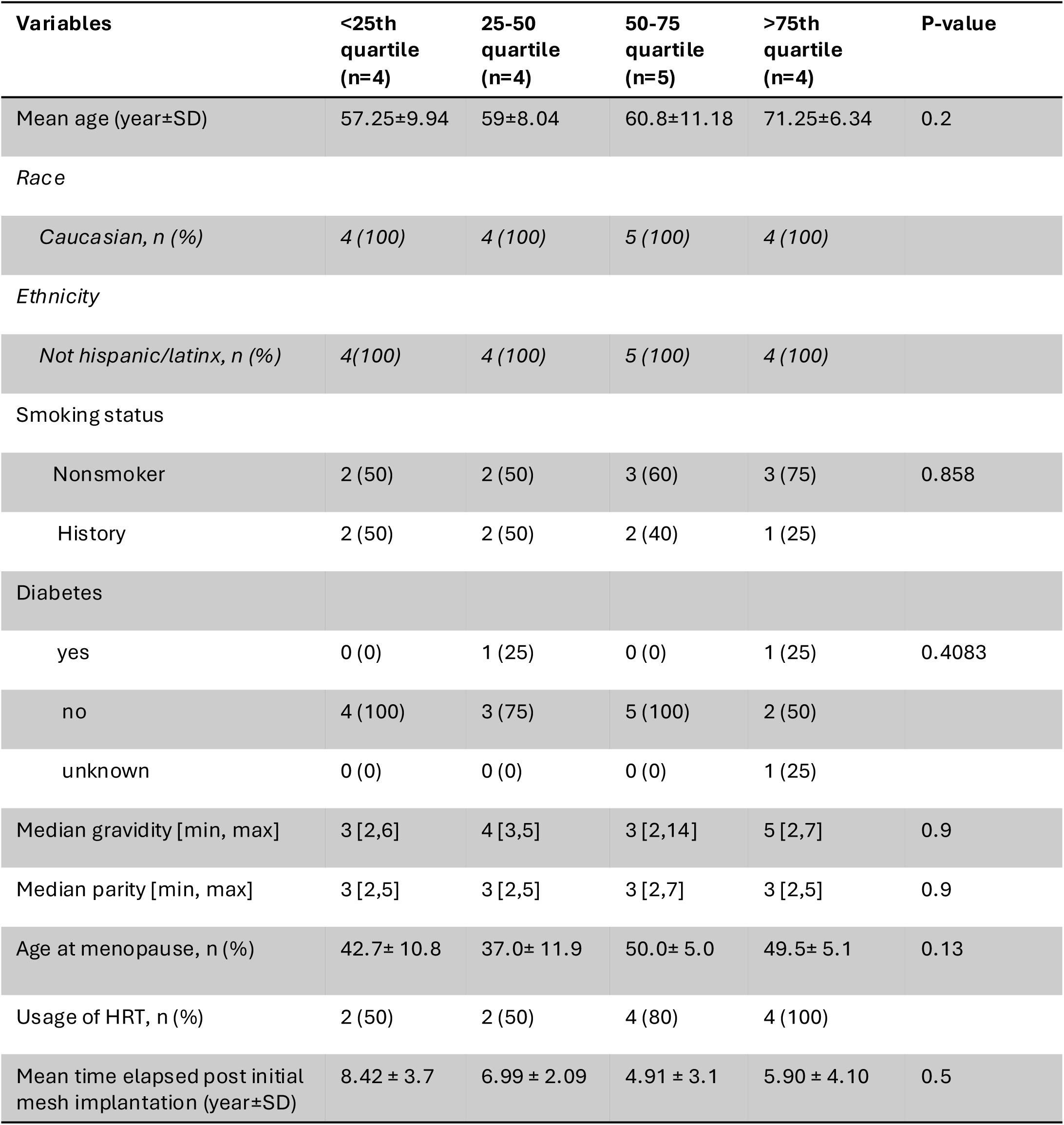
Baseline demographic characteristics of patients stratified by mesh burden quartiles. Values are represented as mean ± standard deviation for continuous variables and n (%) for categorical variables. Mean and Interquartile range is provided for gravidity and parity. P-values were calculated using one-way ANOVA for continuous variables and chi-square tests for categorical variables. *All patients were Caucasian, statistical comparison not applicable. **No patients were current smokers.

### 3.2 Mesh burden quantification

Total mesh area and total adventitial tissue area were quantified, represented as a percent of the total adventitial area occupied by mesh. Samples in the 75^th^-100^th^ percentile had the highest mesh burden (15.2±3.4) compared to the lower quartiles (50-75^th^ 10.2%±1.2% p=0.0085, 25-50^th^ 8.1%±0.5% p=0.008, 0-25^th^ 4.90%±0.9% p<0.0001, **Figure 1A**). Quantifying minimum mesh-to-mesh distance revealed a trend toward lower minimum mesh to mesh distance in the 4th quartile samples as compared to the lower three quartiles (p>0.05, **Figure 1B**).

**Figure 1.**
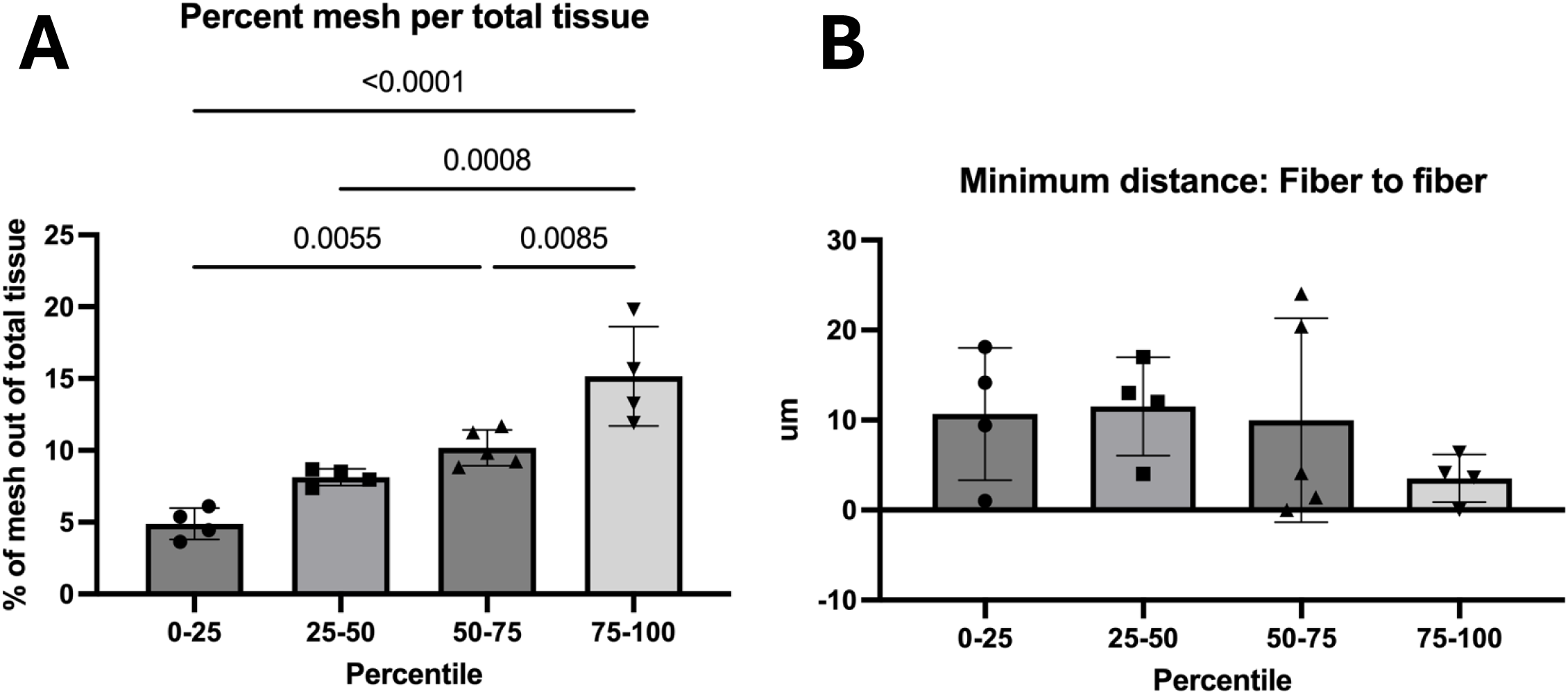
Measured mesh burden from human samples organized into quartiles. Samples removed from women (demographics in Table 1) was quantified histologically as a percent of mesh in the total tissue section. A) Organized into quartiles of increasing mesh burden, this showed significant differences in mesh burden between 0-25^th^ and 50^th^-70^th^ (p=0.005) and 75^th^-100th (p<0.0001), 25-50^th^ and 75-100^th^ (p=0.0008), and 50-75^th^ and 75-100^th^ (p=0.008). B) Minimum distance between mesh fibers was not significantly different between quartiles (p>0.05)

### 3.3 Immunofluorescence cell populations across mesh burden quartiles

Immunofluorescence labelling, imaging, and quantification techniques were used to identify and characterize macrophages (CD68+), fibroblast-to-myofibroblast transitioned (FMT) cells (Vimentin+/αSMA+), and macrophage-to-myofibroblast transitioned (MMT) cells (CD68+/Vimentin+/αSMA+, **Figure 2A**). Upon examining macrophage density (cells/mm^2^ of tissue), there were significantly more macrophages in the highest mesh burden quartile (75^th^-100^th^ 4098.2 ± 564.6 cells/mm^2^) as compared to the first and second quartiles of mesh burden respectively (0-25^th^ 1839.4 ± 262.5 cells/mm^2^, 25-50^th^ 1894.2 ± 600.5 cells/mm^2^, p<0.03, **Figure 2B)**. While there was a steady increase in mean myofibroblast density (vimentin+/αSMA+ with or without CD68) with increasing mesh burden (918.9 ± 389.7 cells/mm^2^, 1238.0 ± 619.2 cells/mm^2^, 1404.7 ± 812.5 cells/mm^2^, 1658.3 ± 755.7 cells/mm^2^, respectively) there was no statistical difference between quartiles (p>0.05, **Figure 2C**). Myofibroblasts were then classified by origin as FMT (vimentin+/αSMA+/CD68-) or MMT (vimentin+/αSMA+/CD68+) and are shown as a percent of total myofibroblasts. In all mesh complication samples, the majority of myofibroblasts were MMT rather than FMT origin across all four mesh burden quartiles (p<0.0001, 75^th^-100=98.4%, 50-75^th^=98.7%, 25-50^th^=95.9%, 0-25^th^=95.7%, **Figure 2E**). MMT (vimentin+/αSMA+/CD68+) density increases with increasing mesh burden, however this was not statistically significant (p>0.05; 0-25^th^ 898.8 ± 398.8 cells/mm^2^, 25^th^-50^th^ 1173.9 ± 566.2 cells/mm^2^, 50^th^-75^th^ 1386.6 ± 808.0 cells/mm^2^, 75^th^-100^th^ 1636.1 ± 765.7 cells/mm^2^, **Figure 2D**).

**Figure 2.**
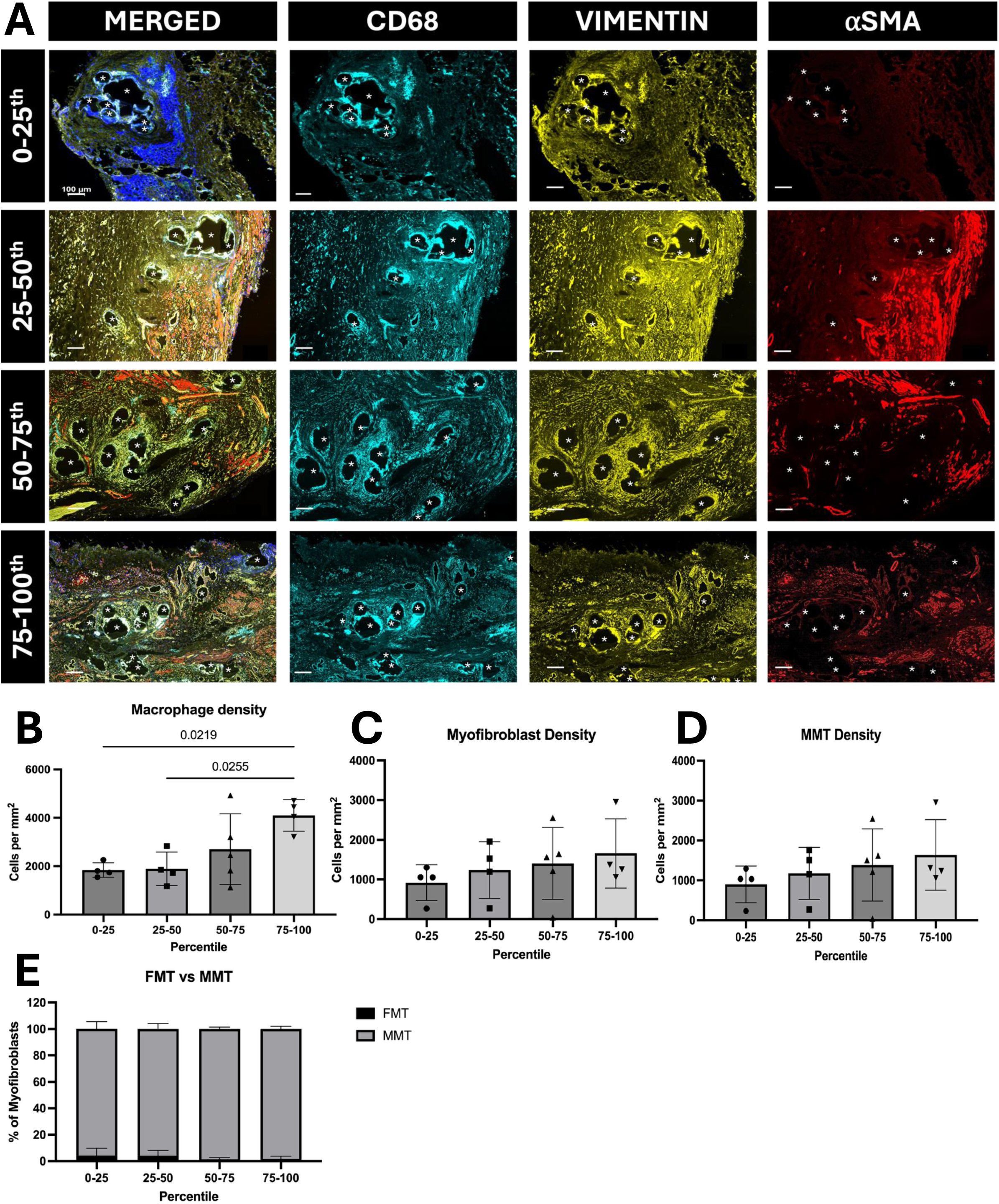
Macrophage density and percentage of myofibroblasts labelling with macrophage marker both increase with mesh burden. A. Representative immunofluorescence image from each quartile showing labelling with CD68 (macrophage marker), vimentin (fibroblast), and *a*SMA (myofibroblast) marker. B. Macrophage density, as measured by CD68+ cells out of total area in the adventitia, increases with increased mesh burden in samples (p=0.02). C. Myofibroblast density, measured by cells labelled vimentin+aSMA+ out of total area, does not significantly increase with mesh burden (p>0.05). D. Macrophage-to-myofibroblast (MMT) density, as measured by myofibroblasts (vimentin+aSMA+) also labelling with CD68+ does not significantly increase with mesh burden (p>0.05). E. Out of all myofibroblasts MMTs (vimentin+aSMA+CD68+) are the main fibroblast phenotype responding to mesh burden, as compared to fibroblast-to-myofibroblast (vimentin+aSMA+) p=0.0001). Scale bars= 100um. * indicated mesh fibers.

Mesh integration into vaginal tissue relies on a spatially organized and distinct response to each fiber typically a foreign body granuloma in which macrophages form the first 2-3 layers closest to the fiber surrounded by a layer of fibroblasts/myofibroblasts and collagen that encapsulates it. To assess how well this organization is maintained in the presence of MMT-derived myofibroblasts, we quantified the distribution of macrophages (CD68+) by distance to the nearest mesh fiber as a measure of the spatial organization of the immune response. In the lowest mesh burden quartile (0-25th percentile), the proportion of macrophages decreased rapidly and significantly with distance from the mesh (19% at 100-200 µm vs 12% at 200-300 µm, p=0.02; and 9% at 300-400 µm, p=0.03; **Figure 3A**), indicating a highly localized macrophage response at the mesh-tissue interface. At higher mesh burden, this distance-dependent pattern was less consistent and diminished at a further distance (25th-50th percentile: 23% at 0-100 µm vs 14% at 100-200 µm, p=0.005 but p>0.05 when 0-100 um was compared to greater distances; 50th-75th percentile: 37% at 0-100 µm vs 8% at 300-400 µm, p=0.01). For the highest quartile of mesh burden, the decrease in macrophages was not clearly significant at any measured distance(45% at 0-100 µm vs 12% at 400-500 µm, p=0.05; all other comparisons p>0.05; **Figure 3A**). This loss of consistent spatial organization may reflect overlapping cytokine gradients from adjacent fibers and is consistent with a less organized host response at high mesh burden.

**Figure 3.**
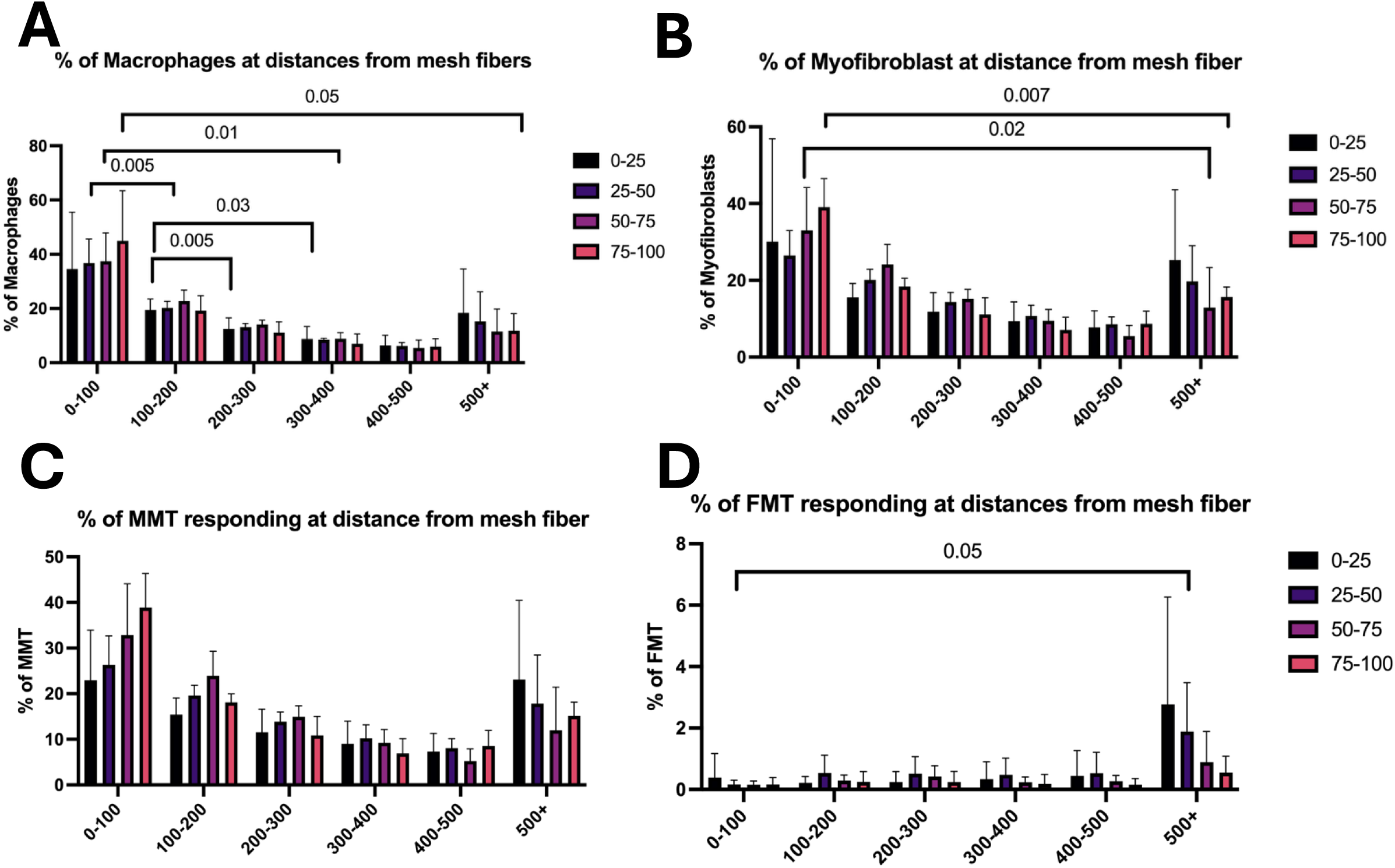
Localization of the cellular response to mesh fibers shows macrophage-to-myofibroblast cells labeling at the mesh-tissue interface, following macrophage foreign body response behavior. Distances were measured as the distance from each cell to nearest mesh fiber. Cells were then organized in 100µm increments from mesh fiber. A. Macrophages (CD68+) show significant localization to the mesh fiber, traditional to macrophage response to a foreign body. B. Myofibroblast (vimentin+/aSMA+) were seen to be evenly distributed from the mesh fiber. C. Macrophage-to-myofibroblasts (vimentin+aSMA+CD68+) were seen more localized to mesh (0-200µm) following a traditional macrophage response. D. Fibroblast-to-myofibroblast (vimentin+aSMA+CD68-) were seen to be farther from the mesh-tissue interface (500+µm) traditional of a myofibroblast responding to a foreign body.

Within the foreign body response, myofibroblasts (vimentin+/αSMA+) are typically located peripheral to the immediate macrophage-rich zone around each fiber, where they deposit matrix and encapsulate the mesh fibers, thereby containing the foreign body response [22, 23]. However, with increasing mesh burden we observed a high number of myofibroblasts localized to the mesh-tissue interface (0-100 µm), with significantly fewer seen at farther (400-500um) distances (0-25^th^ p>0.05, 25^th^-50^th^ p>0.05, 50-75^th^ p=0.02, 75-100^th^ p=0.007, **Figure 3B**). Because these myofibroblasts are localizing to where macrophages traditionally respond, we looked at spatial localization by myofibroblast cell origin. This showed that the localized myofibroblast response was due to MMTs (vimentin+/αSMA+/CD68+) and not FMTs (Vimentin+/αSMA+/CD68-), the latter were observed in the expected location farther from the mesh-tissue interface (500+ µm, p=0.05) (**Figure 3C&D**). There were no differences in cellular localization distances between quartiles (p>0.05).

### 3.4 Trichrome staining for total collagen content

Collagen, the major fibrillar ECM component of the vagina, was assessed by Masson’s trichrome staining and quantified as the percent of total tissue area staining positive for collagen, excluding mesh area (**Figure 4A**). Collagen content did not differ significantly across mesh burden quartiles (p>0.05; **Figure 4B**). However, when normalized to mesh area, the collagen-to-mesh ratio was significantly higher in the 0-25th percentile than in the 25th-50th (p=0.0005), 50th-75th (p=0.005), and 75th-100th (p=0.0002) percentiles (**Figure 4C**), indicating that at higher mesh burden a similar amount of collagen supports a greater amount of mesh. This lower collagen-to-mesh ratio is consistent with stress shielding, a proposed mechanism for mesh exposure, in which load-bearing is shifted to the mesh and the surrounding tissue is under-stimulated [22, 23].

**Figure 4.**
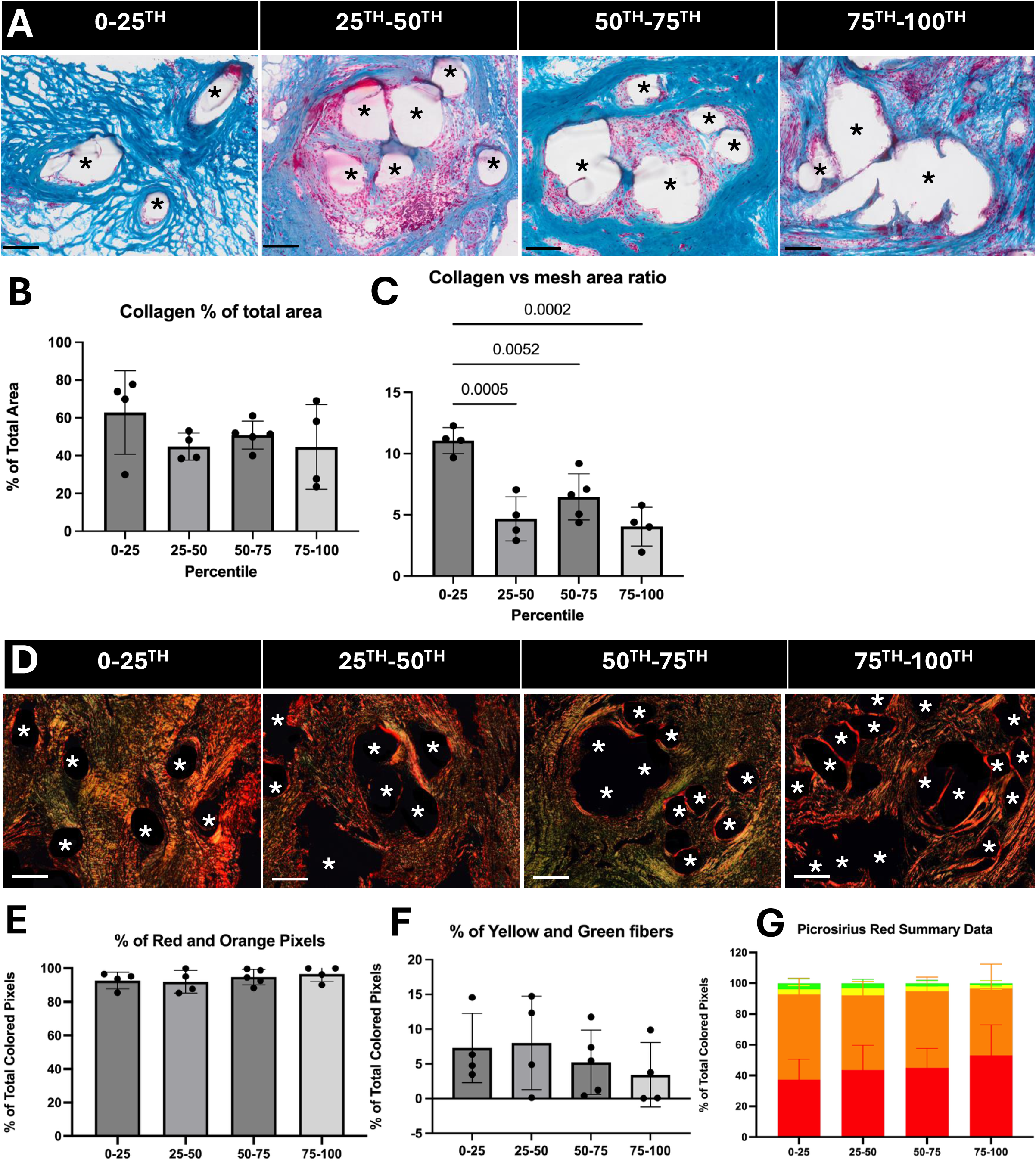
Collagen deposition decreases with mesh burden, with an increase in mature/fibrotic collagen deposition. **A.** Massons Trichrome images were stained and brightfield imaged. Scale bar is 100um. * indicates mesh fibers. B. Collagen as a % of total adventitial area was not different across groups (p>0.05). C. Collagen area vs mesh area ratio decreased with increasing mesh burden (p=0.0002). D. Picrosirius red staining imaged under polarized light. Scale bar is 100um. * indicated mesh fibers. E. % of Red and Orange pixels out of all color pixels showed no difference across groups (p>0.05). F. % of yellow and green pixels out of all color pixels showed no difference between mesh groups (p>0.05). G. Within groups, there was more red than yellow or green in groups with higher mesh burden (50-75, p=0.01, 75-100, p=0.05).

### 3.5 Picrosirius red stain

Picrosirius red stained slides imaged under polarized light was used to assess collagen fiber organization with thicker, more mature collagen fibers appearing red/orange, and thinner, newly deposited fibers appearing as yellow or green (**Figure 4D**). The proportions of red/orange (p>0.05; **Figure 4E**) and yellow/green (p>0.05; **Figure 4F**) collagen did not differ across mesh burden quartiles. Within each quartile, however, red/orange collagen predominated over yellow and green collagen (0-25th: p=0.005 and p=0.007; 25th-50th: p=0.001 and p=0.003; 50th-75th: p=0.0006 and p=0.0008; 75th-100th: p=0.03 and p=0.03). Red collagen was also more abundant than yellow or green collagen in the 50th-75th (p=0.01, p=0.008) and 75th-100th (p=0.05, p=0.04) quartiles; **Figure 4G**. Together with the trichrome data, these findings indicate that collagen was predominantly composed of thicker, more mature fibers regardless of mesh burden, with no evidence that the proportion of newly deposited collagen differed across quartiles.

### 3.6 Proteomic analysis

To more comprehensively evaluate the biological response to mesh burden across groups, we assessed tissue-level differences in immune cell and remodeling-associated markers in a proteomic atlas of patients undergoing mesh excision [5]. The cohort consisted of samples collected from women who had mesh removed as a result of exposure (N=10), pain (N=10), or had a well-integrated (N=10) mesh and desired mesh removal (Magee Women’s Mesh and Pelvic Floor Tissue Biorepository). Participant age was similar across mesh-exposure (52 ± 7 years), mesh-pain (49 ± 14 years), and well-integrated (62 ± 13 years) groups. However, the control group (42 ± 6 years) was significantly younger than the mesh-exposure group (p<0.05). The patients in the proteomic cohort are significantly younger (50.8 ± 12 years) than the cohort described in the above analysis (62 ± 10 years, p=0.002). There were no differences in mean body mass index (BMI) was 28 ± 6 kg/m^2^, and median duration of implantation was 6 ± 5 years between the proteomic cohort and mesh exposure cohort described above, or between the groups (p>0.05). The proteomics cohort was predominantly white, with one participant in the mesh-exposure group identifying as Hispanic/Latinx.

A predefined list of 40 proteins of interest involved in upregulation of macrophage, myofibroblast, and ECM deposition was cross-referenced against the proteomics dataset using 70 possible protein identifiers, resulting in the identification of 19 proteins. Mannose Receptor C - type-1 (MRC1, an M2-like macrophage marker) and pan-macrophage marker (CD68) demonstrated 7-8 log2fold increased expression in exposure compared to control vaginal biopsies (p=0.005, p=0.0007, **Figure 5A**). The fibroblast-associated proteins VIM, LAP3, TNC, and FAP were found to 1-2 log2fold increased in exposure (adjusted p value ≤ 0.02, **Figure 5A**). TGFB1I1 was found to be negatively differentially expressed 2 log2fold in all complication groups, significantly in exposure (p=0.05, **Figure 5A**). The extracellular matrix-associated proteins, MMP9, was found to be significantly upregulated in exposure as compared to well integrated (p=0.001), as well as FN1 (p=0.012). COL1A2, COL1A1, COL3A1, MMP2, and CD44 were not found to have a significant difference between groups (p>0.05, **Figure 5A**).

**Figure 5.**
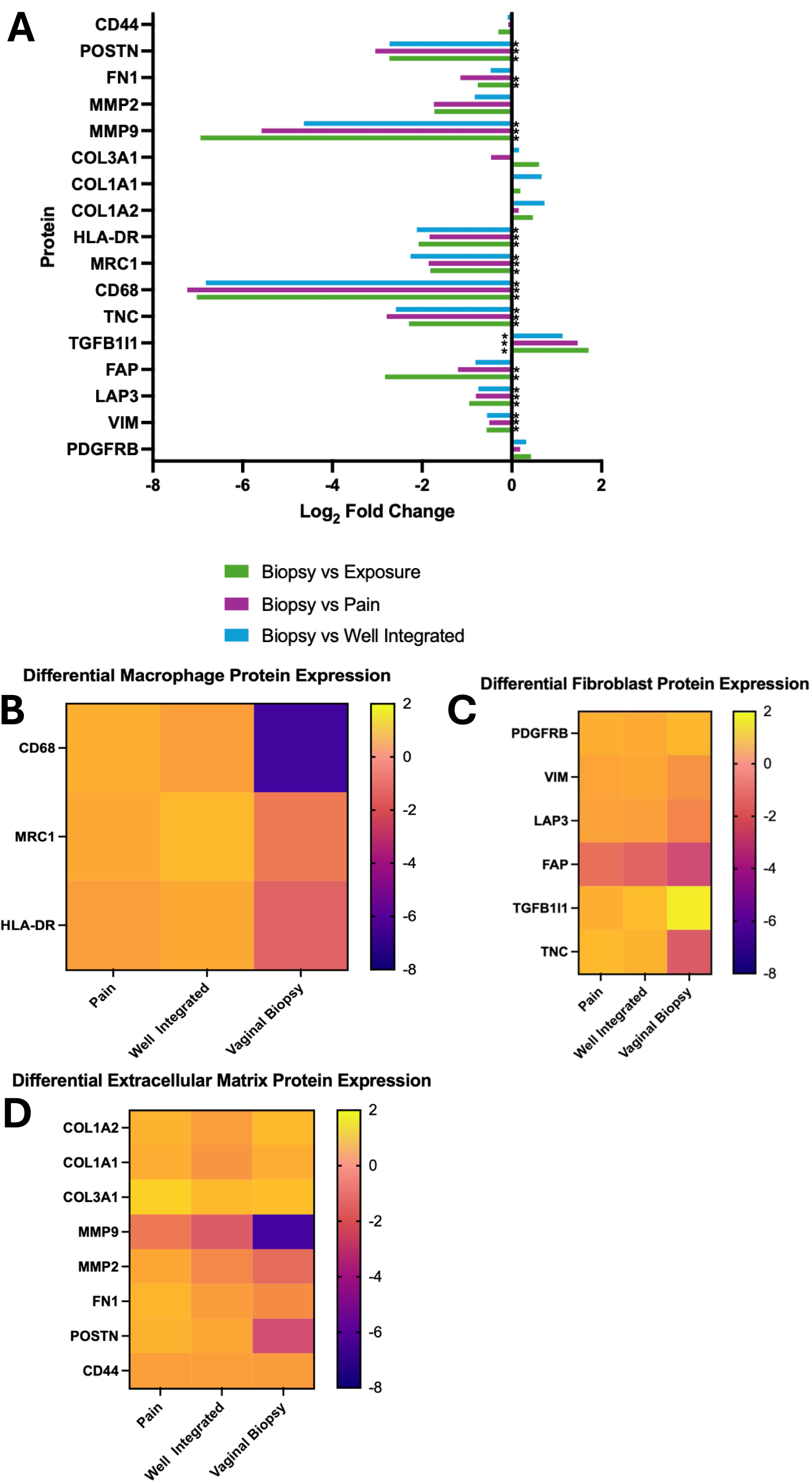
Proteomic analysis shows increased macrophage, collagen 1, and matrix degrading enzymes in exposure. Differential protein expression was measured from exposure, pain, well-integrated, and vaginal biopsies from a tissue biorepository. Differential exposure is expressed by log2fold change as a comparison from A. Vaginal biopsy as a measure of the effect of mesh placement, and B. Exposure vs pain, well-integrated, and biopsy to measure the effect of mesh complications of macrophage, fibroblast, and extracellular matrix proteins.

We then compared differential protein expression in exposure samples compared to the other mesh groups, expressed as log2fold change. Macrophage associated protein expression markers - MRC-1 and CD68-were 2 log2fold higher in exposure compared to pain and well-integrated, and 8 log2fold more than vaginal no mesh control (adjusted p-value ≤ 0.001, **Figure 5B**). Fibroblast differential protein, Vimentin, LAP3-expression was found to be 1 log2fold increased in exposure compared to pain, well-integrated, and vaginal control biopsy (adjusted p value ≤ 0.02, **Figure 5C**). TGFB1l1, activated in response to mechanical signals, was upregulated in well-integrated and pain, and increasingly upregulated to 2 log2fold in exposure as compared to vaginal control biopsy (adjusted p-value <0.0001, **Figure 5C**). Tenascin (TNC) expression was 4 log2fold upregulated in exposure compared to pain, well-integrated, and biopsy (adjusted p value ≤ 0.02, **Figure 5C**). Extracellular matrix proteins-COL1A2, COL1A1- were upregulated in exposure 1 log2fold change compared to pain (adjusted p value ≤ 0.02) but not well-integrated or biopsy (adjusted p>0.05, **Figure 5D**). Collagen 3 (COL3A1) was not differentially expressed across groups (adjusted p>0.05, **Figure 5D**). Matrix-degrading protease MMP9 was significantly upregulated 4-6 log2fold in exposure compared to pain, well-integrated, and vaginal control biopsy (adjusted p value ≤ 0.04). MMP2 is upregulated 1 log2fold in exposure compared to other outcomes, but not significantly (adjusted p>0.05, **Figure 5D**).

The proteomic data demonstrate mesh exposure is associated with increased macrophage-associated proteins, selective upregulation of fibroblast proteins, and enhanced extracellular matrix remodeling, specifically collagen 1 associated with fibrotic matrix deposition. This supports the targeted histologic findings presented here demonstrating macrophage activation and transition towards fibroblast-like features, a possible mechanism for clinical complications.

## Discussion

In this study, we examined the macrophage and myofibroblast response to polypropylene mesh in tissue from women with mesh complications, stratified by mesh burden. Prior work in a non-human primate model showed that mesh deformation, which increases mesh burden, is associated with a larger macrophage response, a less organized cellular architecture, and a higher proportion of myofibroblasts co-expressing CD68 [14, 19]; to our knowledge, this is the first analysis of the myofibroblast response in mesh explanted from women. Macrophage density increased with mesh burden, and the spatial organization of macrophages around individual fibers was less consistent at higher burden. Myofibroblast density did not differ across quartiles, but nearly all myofibroblasts in every quartile co-expressed CD68 and were concentrated near mesh fibers, similar to the highest-burden primate conditions [19]. This is consistent with macrophage-to-myofibroblast transition (MMT), although marker co-expression alone cannot establish this origin, and because all samples were from complications we cannot determine whether it is specific to complications.

Collagen as a percent of tissue area did not differ with burden, but collagen per unit of mesh was lower at higher burden, and collagen was predominantly a mature subtype in all quartiles. In a separate, younger proteomic cohort, CD68, MRC1, and MMP9 were higher in exposure than in pain, well-integrated, and non-mesh vaginal tissue, consistent with the elevated MMP9 observed in the highest mesh burden groups in nonhuman primate studies [19]. Together, these findings suggest an actively remodeling, macrophage-rich environment in exposure and generate the hypothesis that MMT contributes to mesh complications. Increase in mesh burden across groups was found not to be necessarily driven by decreased distance between fibers but potentially tissue degradation as a response to mesh. Although mesh contraction and pore collapse are known risk factor for mesh complications [19, 24, 25], this corroborates that in patients with exposure, tissue degradation is also contributing to effective mesh burden.

As previously defined, macrophages are a key player in the host response to implanted mesh for prolapse repair [8, 14, 21]. Here we add to this knowledge by defining the macrophage-derived myofibroblast as >95% of the myofibroblasts found at the mesh-tissue interface in exposure complications. This is supported by CD68+ coexpression with vimentin and alpha-smooth muscle actin and spatial localization consistent with a macrophage response. MMT cells have been identified in other organ systems in fibrotic environments [16, 26, 27]. This transdifferentiation has been seen in renal allograft rejection, cardiac fibrosis, and in promoting cancer progression in lung cancer. In these studies, MMT cells were observed to take on the role of resident fibroblasts, and promote fibrotic tissue organization [26, 28, 29]. Previous work has identified the role of MMTs in a non-human primate prolapse-repair model, showing MMTs as the primary myofibroblast responding to implanted mesh, with resulting dysregulated matrix deposition [19].

Further supporting the role of myofibroblasts in mesh complications, we observed increased thickened/mature collagen in samples with high mesh burden, despite decreased total collagen, suggesting a fibrotic response to mesh burden. Interestingly, this fibrotic response was seen in samples removed from patients due to exposure. This supports a highly localized response to mesh, with regions of fibrotic deposition and regions of vaginal degradation/thinning -to the point of exposure-in the same patient sample.

Probing of our published proteomic atlas provided valuable insights into differences between complicated (pain, exposure) and uncomplicated (well-integrated) vaginal mesh integration and healing. The data further supported the role of CD206^+^ macrophages (MRC1 and CD68), matrix degradation (MMPs), and fibroblast involvement (Vim, LAP, FAP) in all mesh groups compared to vaginal biopsy. Interestingly, we saw a significant upregulation of TGF-B1I1 with the introduction of mesh compared to the vaginal biopsy. TGFB1I1 is an inducible protein that is activated through the TGF-B signaling pathway to prompt cell adhesion, respond to physical strain, and promote healing [17, 30]. The TGF-B signaling pathway has been established as the main differentiation pathway of MMT trans differentiation [16, 17, 26, 28, 31]. Therefore, these data suggest that the pathway prompting MMT differentiation is upregulated when mesh is present, even in well-integrated mesh.

In comparing mesh complications to well-integrated mesh, the most significant differences were between exposure and vaginal biopsy (MMP9, CD68). Other derivations in expression from vaginal biopsy in macrophage, extracellular matrix, or fibroblast regulatory proteins did not differ between mesh groups. This further illustrates that while these pathways, and specifically MMT differentiation, are activated as a response to mesh, further studies are needed to determine the mechanistic “tipping-point” as to when these cells result in maladaptive collagen deposition and fibrosis.

### Limitations and future directions

This studies provides, for the first time, and insight into the role of the MMT response in women who have had their mesh removed due to complications. These samples allowed us to investigate the role of the MMT response in clinically relevant samples, to inform future work and potential device development. With our study, and the proteomic dataset, we were able to investigate the role of MMT activation pathways in complicated and uncomplicated samples.

Because samples are obtained from mesh excision surgeries, tissue is primarily adventitia rather than full thickness vagina which is a difference compared to full thickness vagina obtained in animal models. Further work in animal models could focus on key mechanisms responsible for the upregulation of the MMT response as a key regulator of complications and the specific factors leading to the disorganized host encapsulation response which ultimately is the underlying mechanism leading to mesh exposure. The current study provides additional evidence that mesh burden drives unfavorable downstream outcomes. Thus, defining all the contributors to mesh deformation (device, surgical, patient) so as to maintain mesh burden at a low level after suturing, tensioning and loading is a critical next step to preventing mesh complications.

## Conclusion

In this study we found that in women with mesh exposure and high mesh burden, the macrophage response is heightened, with a significant increase in macrophage-to-myofibroblast (MMT) transdifferentiated cells. We found that with higher mesh burden, and a higher percentage of MMTs, there was decreased collagen density but more mature collagen, a potential mechanism for complications seen clinically. This was further supported by significant differential expression of key proteins in the MMT response (CD68, TGFB1I1, Vim) in mesh exposure samples To our knowledge, this is the first time the MMT response has been described in human samples as a response to polypropylene mesh implanted for prolapse repair and has the potential to inform future studies in vaginal immunology and wound healing as well as work to design improved vaginal mesh materials and decrease mesh complications.

## Data Availability

The data that support the findings of this study are available from the corresponding author P.A.M., upon request.

## Acknowledgements

The authors are grateful for the financial support for this work provided by the National Institutes of Health R01 HD083383 grant funds to PAM. MAT is supported by the National Center for Advancing Translational Science of the National Institutes of Health under award number TLTR001858. AA and KK is supported by NIH/ORWH Building Interdisciplinary Research Careers in Women’s Health (BIRCWH) NIH K12AR084218. The content is solely the responsibility of the authors and does not necessarily represent the official views of the National Institutes of Health.

